# Post-transcriptional regulation of adult CNS axonal regeneration by Cpeb1

**DOI:** 10.1101/125096

**Authors:** Wilson Pak-Kin Lou, Alvaro Mateos, Marta Koch, Stefan Klussman, Chao Yang, Na Lu, Stefanie Limpert, Manuel Göpferich, Marlen Zschaetzsch, Carlos Maillo, Elena Senis, Dirk Grimm, Raul Mendez, Kai Liu, Bassem A. Hassan, Ana Martin-Villalba

## Abstract

Adult mammalian CNS neurons are unable to regenerate following axonal injury, leading to permanent functional impairments. Yet, the reasons underlying this regeneration failure are not fully understood. Here, we study the transcriptome and proteome shortly after spinal cord injury. Profiling of the total and ribosome-bound RNA in injured and naïve spinal cords identify a substantial post-transcriptional regulation of gene expression. In particular, transcripts associated with nervous system development were downregulated in the total RNA-fraction while remaining stably loaded onto ribosomes. Interestingly, motif association analysis of post-transcriptionally regulated transcripts identified the cytoplasmic polyadenylation element (CPE) as enriched in a subset of these transcripts that was more resistant to injury-induced reduction at transcriptome level. Modulation of these transcripts by overexpression of the CPE binding protein, Cpeb1, in mouse and Drosophila CNS neurons promoted axonal regeneration following injury. Our study uncovers a global conserved post-transcriptional mechanism enhancing regeneration of injured CNS axons.

## INTRODUCTION

Axons of the adult mammalian CNS have a very limited regenerative capacity following injury. Extrinsic factors with inhibitory effect on regeneration have received a great deal of attention, and many molecules have been identified (Filbin, 2006). However, the limited success obtained by strategies based on neutralizing inhibiting signals within the immediate environment of an injured axon(Côté et al., 2011; Young, 2014), has recently turned the attention to cell intrinsic factors involved in positive and negative modulation of the regenerative response (Liu et al., 2011; Mar et al., 2014).

Early in 1913 Santiago Ramón y Cajal already described that besides the weak and sterile end of axotomized axons set to degenerate, there are active axonal ends, capable of sprouting, for which he termed “bud” or “club of growth”, due to their analogy to the growth cones of embryonic axons (Cajal et al., 1991). Interestingly, formation of sprouts at the axonal tip of axotomized dorsal root ganglion (DRG) neurons is accompanied by increased expression of regeneration-associated genes (Ylera et al., 2009). Successful regrowth of central DRG axons as induced by a preconditioning peripheral lesion requires the assembly of these regenerating terminal bulbs that are observed 5-7 hours following injury (Ylera et al., 2009). Processes like membrane sealing, regulation of proteolytic processes, RNA stability and local translation are major determinants of successful assembly of a regenerating axonal terminal. Therefore, injured CNS axons do attempt to regrow in the early post-injury phases but ultimately fail to do so. Consistent with this it has been shown that intrinsic pro-regenerative response has to be stimulated before the onset of overt inflammatory response and scar formation (Bradke et al., 2012). Finally, and despite considerable phylogenetic distance between Drosophila and mouse, many phenomena and mechanisms of CNS development and regeneration are conserved between the two species (Hoopfer et al., 2006; MacDonald et al., 2006; Song et al., 2012), including the transient abortive regeneration after CNS injury(Ayaz et al., 2008). This allows application of results from fly genetics to speed up finding and validation of candidate regulator genes for regeneration in mouse models

Post-transcriptional regulation of gene expression is of particular importance in neurons, as it allows enhanced spatial and temporal control in locations far removed from the cell soma, such as synapses and axonal growth cones (Holt and Schuman, 2013; Jung et al., 2012). Indeed, axonal regeneration in peripheral sensory neurons involves post-transcriptional regulation, with localized translation of specific transcripts at the tip of the injured axon (Hanz et al., 2003; Perlson et al., 2005; Perry et al., 2012; Yan et al., 2009; Yudin et al., 2008). Interestingly, stability and local translation at the injured axon of some of these regeneration-regulating transcripts is mediated via the 3’UTR of the transcripts. For example, in *C. elegans* adult touch neurons, axotomy induces activation of DLK1 which promotes stabilization and local translation of CEBP-1 by modifying its 3’UTR, and leads to robust regeneration (Yan et al., 2009). 3’ UTR-localized elements are known to nucleate the formation of ribonucleoparticle complexes by binding miRNAs and/or RNA-binding proteins, which in turn, through transport/compartmentalization and regulation of mRNA poly(A) tail length, controls translation and RNA stability (Weill et al., 2012). Some of these motifs include cytoplasmic polyadenylation element (CPE), Pumilio binding element (PBE), Musashi binding element (MBE), and AU-rich elements (AREs) (Charlesworth et al., 2013; von Roretz et al.). CPE is best known for its role in cytoplasmic polyadenylation, where it regulates the length of the poly(A) tail of the transcripts that contain it and thus its translation(Ivshina et al., 2014; Weill et al., 2012). CPE-mediated regulation can be complex, as it could act in conjunction with other cis elements such as PBE, MBE, AREs or miRNA binding elements whereby the number, arrangement and distance between motifs and availability of their regulators such as kinases or RBPs have varying effects on transport, stability or translation of the transcript (Piqué et al., 2008; Weill et al., 2012). In the nervous system, Cpeb1 is known to transport transcripts to postsynaptic densities in dendrites, where it promotes their translation upon synaptic activity (Huang et al., 2003; Udagawa et al., 2012; Wu et al., 1998).

We reasoned that the early transient regenerative response might offer key insights into the intrinsic molecular mechanisms that need to be activated for successful regeneration. To this end, we profiled total and polysome-bound RNA of sham operated (naïve) and acutely-injured spinal cords 9h following injury –a time when infiltration of the spinal cord by inflammatory cells has just started (Letellier et al., 2010). We find that changes in the two RNA fractions are highly uncoupled, and that uncoupled genes could successfully predict axonal growth regulators in a *Drosophila* model. Presence of the CPE motif in some of these transcripts correlated with resistance to injury-induced decrease of transcript availability. Finally, we show that in a drosophila axonal-injury model and a mouse model of optic nerve crush-injury, axonal regeneration is enhanced upon Cpeb1 overexpression. This study uncovers a highly conserved global switch in neurons that increases availability of regeneration-associated-RNA transcripts and thereby enables axonal regeneration following axotomy.

## RESULTS

### Transcriptome and translatome responses to spinal cord injury are extensively uncoupled

To study post-transcriptional regulation in the spinal cord during the abortive regeneration phase following injury, we performed simultaneous profiling of the transcriptome and translatome from injured and naïve mice via translation state array analysis (TSAA). Total and polysome-bound RNAs were extracted from spinal cord tissues of injured and naïve mice 9 hours after operation (Fig. 1A). This time point was chosen to be within the transient regenerative phase after spinal cord injury, thereby allowing sufficient time for transcriptional and translation changes to take place, under limited immune response conditions (Gadani et al., 2015; Hausmann, 2003; Schwanhäusser et al., 2011; Trivedi et al., 2006). Spinal cord tissue surrounding the injury site (2.5 cm distal/proximal) was used for RNA profiling to gain insights into the processes taking place in the proximal and distal axonal ends affected by the injury. Polysome-bound transcripts were isolated via sucrose gradient fractionation, and fractions containing RNA bound to two or more ribosomes were collected. Samples were subsequently hybridized onto Affymetrix microarrays. Total and polysome-bound RNAs were normalized separately, as polysome-bound RNA is a subset of total RNA and standard microarray normalization methods that assume equality of distributions of total intensity between arrays could not be used. Notably, correlation plots between arrays shows low variability between replicates, and assessment of expression changes by quantitative real-time PCR (qPCR) largely validated the expression changes obtained from microarray analysis (Fig. S1A-B).

**Figure 1:**
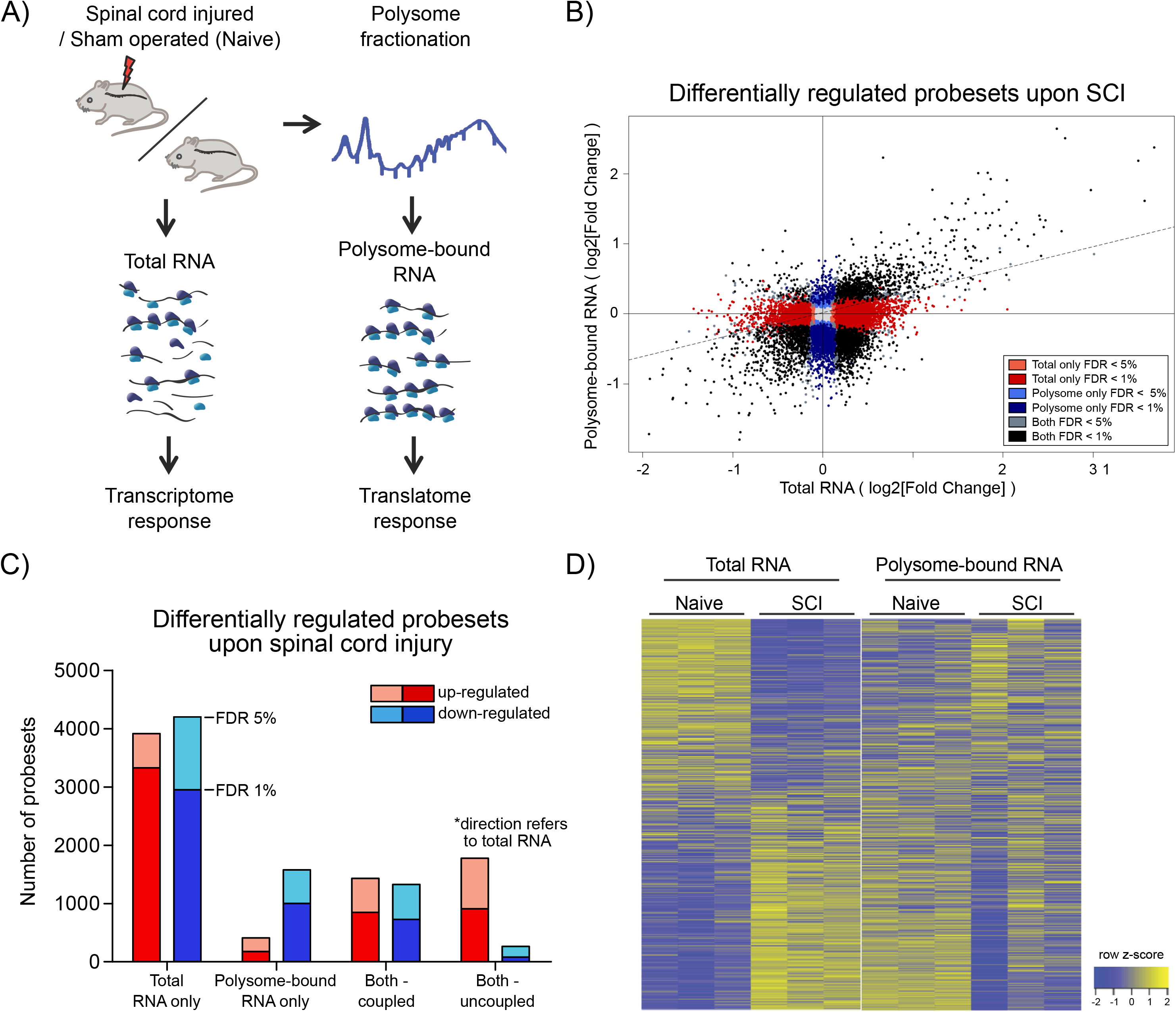
Wide-spread uncoupling of transcriptional and translational responses following spinal cord injury. A) Experimental scheme of simultaneous profile of the transcriptome and translatome. Total or polysome-bound RNA fractions were extracted from naive or injured spinal cords and analyzed by RNA microarray. 3 mice were used as biological replicate within each experimental group. B) Scatter plot representation of fold changes of all probesets in total and polysome-bound RNA upon spinal cord injury. C) Number of differentially expressed probesets. D) Heat map representation of probeset expression as z-score in each sample. Each column represents one biological replicate and each row represents expression of one gene across columns. FDR: Benjamini-Hochberg false discover rate.

Since the chosen tissue not only contains the injured neuronal processes, but also other cellular subtypes, we first analyzed whether the injury would have a major impact on cellular tissue composition at this early time point. To this end we compared the intensities of probesets mapped to marker genes for motor neurons and other local neurons, oligodendrocytes, microglia, precursor cells present in the central canal, and blood-borne cells (see Table S2). The analysis revealed a high correlation between expression patterns of naïve and injured spinal cord (0.97 and 0.99 Pearson correlation for total and polysome fraction, respectively), indicating the absence of major changes in tissue composition upon injury (Fig. S1C). By contrast, a high proportion of probesets in both total and polysome-bound fractions exhibited significant changes upon injury (Fig. 1B). Importantly, for many differentially expressed probesets, the changes in total and polysome-bound RNA do not correlate (Fig. 1B-D, Table S1). A large proportion of changes occur only in the total RNA fraction with no corresponding changes in the polysome-bound RNA. This agrees with previous observations in which stress conditions trigger a general shut down of translation to maximize cell survival (Park et al., 2008; Yamasaki and Anderson, 2008). Differentially regulated genes in the total RNA fraction are similarly distributed between up- and down-regulation (1B-C). The polysome-bound RNA fraction showed fewer differentially regulated genes than the total RNA fraction, with most of those being down-regulated (Fig. 1B-C). The difference in numbers of differentially regulated genes between the total and polysome-bound fractions indicates that the translational response to injury is highly uncoupled from RNA availability. In addition, many genes displayed opposing directions of regulation, suggesting considerable influence of post-transcriptional regulation (Fig. 1D).

### Uncoupled genes are functionally clustered and regulate neuronal regeneration

To assess the functional role of the observed uncoupling effect, Gene Ontology (GO) (Ashburner et al., 2011) enrichment analysis was performed and visualized using Cytoscape (Shannon et al., 2003). Enrichment of up- and down-regulated genes was represented as red and blue nodes respectively. In the total-RNA fractions, transcript availability of genes related to translation, RNA processing, protein catabolic processes and protein transport was increased upon injury, but decreased for genes related to CNS development (Fig. 2A and Table S3). Excitingly, in the polysomal-bound RNA fraction, injury increased ribosome-loading of genes related to regulation of CNS development, as well as cell death, transcription, RNA processing and immune response (Fig. 2B and Table S3). Notably, there were no significantly under-represented categories in the polysome fraction, indicating that the decrease in translation after injury is a general effect and neither directed nor functionally clustered.

**Figure 2:**
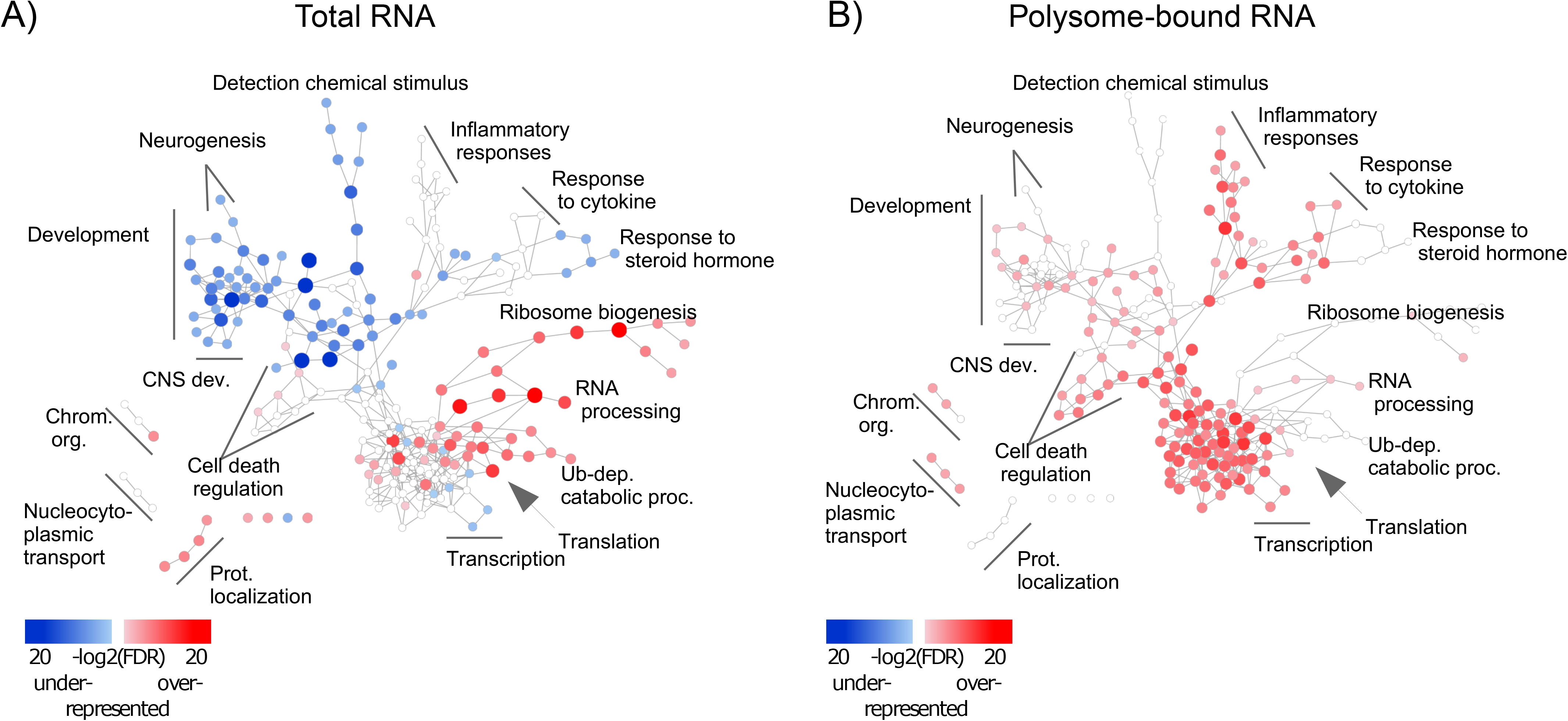
Injury response from total and polysome-bound RNAs is functionally clustered. Gene Ontology enrichment of differentially regulated genes in A) total and B) polysome-bound RNA fractions, represented as a network of GO categories. Enrichment analysis performed as up-regulated genes against all differentially regulated genes. Under-representation is equivalent to an enrichment of down-regulated genes. Colour intensity and size of the node represent significance by FDR. Only significantly enriched GO categories (FDR < 1e-4) are shown.

Different trends of enrichment were observed for many GO categories between the total and polysome-bound fractions, suggesting that uncoupling serves specific functional purposes. Of particular note, categories related to CNS development, which are decreased in the total RNA fraction, remain stable or enriched in ribosomal-loaded transcripts. This might explain the temporary regeneration observed following injury, which is absent at later stages of the injury response, as existing transcripts from regenerative genes continue to be translated at first, but are not replaced upon eventual degradation. To investigate if transcripts exhibiting this highly uncoupled behavior affect axonal growth, we turned to *Drosophila*. Using the UAS-Gal 4 (Brand and Perrimon, 1993), we expressed each of a total of 38 candidate uncoupled genes - for which a fly homologue exists and a UAS line was available - in a fly CNS neuronal population called the small ventral lateral neurons (sLNvs). Most of the genes tested have reduced transcript availability but stable ribosomal loading in both naïve and injured spinal cord (Table S1). We find that 19 (50%) of tested candidates influenced the developmental growth of the sLNv axonal projection: 12 increased outgrowth, while 7 resulted in shorter sLNv projections (Table 1).

**Table 1:**
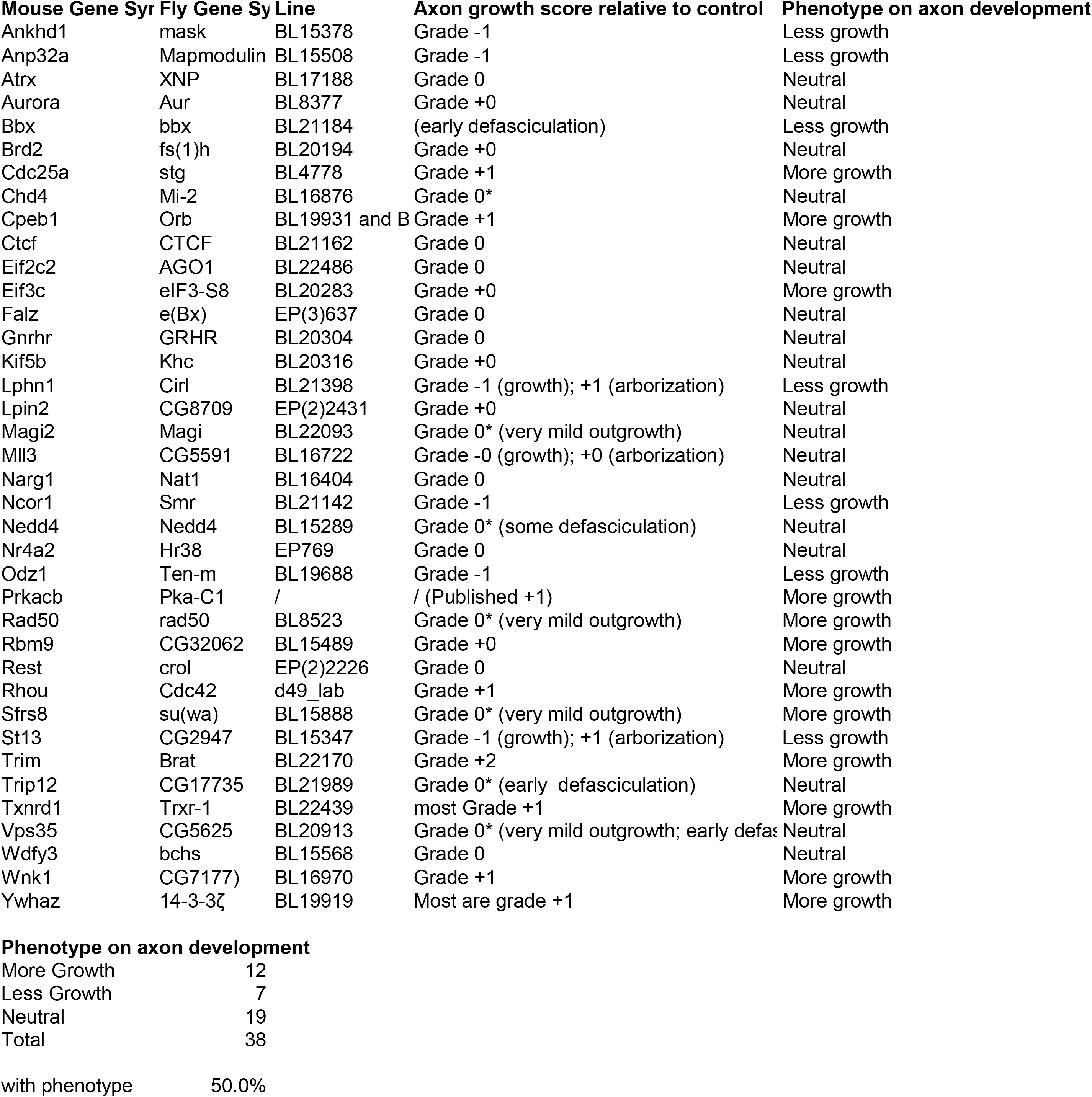
List of genes tested in Drosophila axonal outgrowth screening, the observed phenotypes and the corresponding expression changes in microarray

### Association of 3’UTR motifs with attenuated decrease in transcriptome following spinal cord injury

Next, we asked whether there are common features shared among the uncoupled transcripts, especially since the *Drosophila* experiments suggest that such transcripts are enriched for genes modulating axonal growth. To address this question, we examined the presence of common RNA features. It has been shown that the 3’UTR harbours a myriad of regulatory motifs that regulates stability, intracellular localization and translation of its RNA host (Moore, 2005; Szostak and Gebauer, 2013). Many important neuronal mRNAs such as Map2, Bdnf, β-actin and Gap43 are regulated via their 3’UTRs (Blichenberg et al., 1999; Donnelly et al., 2011; Lau et al., 2010). This prompted us to investigate the association of 3’UTR motifs in a given transcript with its expression upon injury. Since this has to be performed on the transcript level, only probe sets mapping to a unique transcript were used. We investigated the presence of cytoplasmic polyadenylation element (CPE), Pumilio binding element (PBE), Musashi binding element (MBE), Hex (hexanucleotide involved in polyadenylation) and AU-rich elements (AREs) (Table S4). To investigate differences in expression changes upon injury relative to motif-free transcripts, we plotted the density curves showing the probability of a data point to have a given log2-fold change, thus reflecting the pattern of distribution of expression of the set of transcripts of interest (Fig. 3).

**Figure 3:**
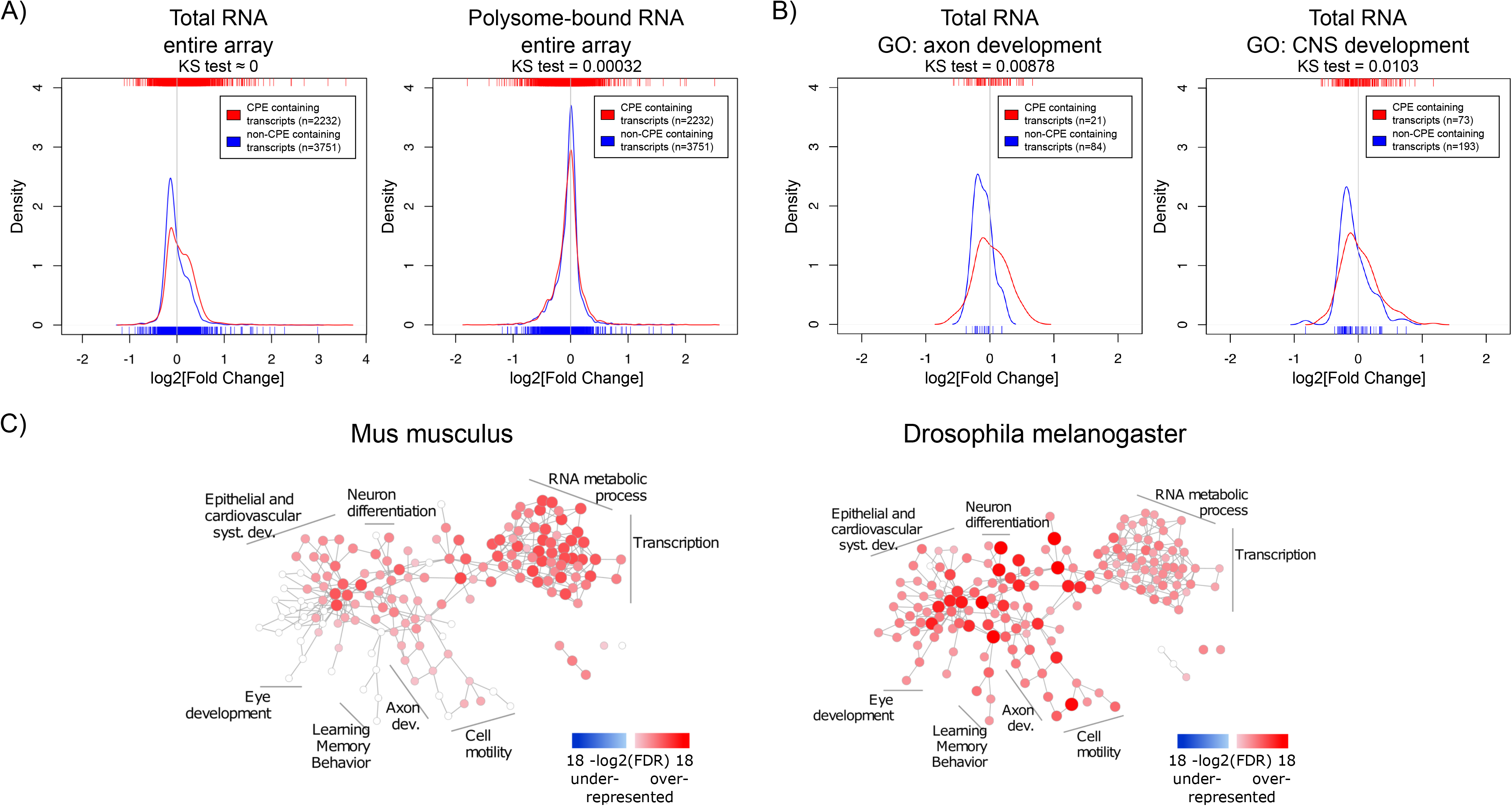
The CPE-motif is enriched in transcripts showing higher positive expression changes following spinal cord injury in the total RNA fraction and whole-genome wide in transcripts related to developmental processes. A-B) Density curves showing the distribution of fold changes (FC) in expression upon injury of CPE-containing and CPE-free transcripts. A) Distribution of FC at the total and polysome-bound RNA level. B) Density curve of FC at the total RNA level of genes associated with GO categories of axon and CNS development. Ticks on top and below the plots represent values of log2 (fold change) of individual transcripts. Distributions were compared with Kolmogorov-Smirnov test. C) Enrichment of CPE-containing genes in mouse and Drosophila genomes represented as network of GO categories. Intensity of colour and size of node represent level of significance. Only GO categories significant in any of the genomes (FDR < 1e-5) are shown.

This analysis revealed that transcripts possessing CPE are associated with resistance to injury-induced down-regulation, as compared to transcripts devoid of CPE (Fig. 3A). Notably, this association was also seen for genes related to axon and CNS development GO categories (Fig. 3B). In contrast, presence or absence of CPE did not influence injury-induced changes on the level of ribosomal-loading (Fig. 3A and S2A). Likewise, RNA transcripts possessing PBE, MBE, Hex and AREs were also associated with resistance to injury-induced down-regulation in the total but not in the polysomal RNA fractions (Fig. S2B-C, S3A-B). CPE and AREs were also found to co-occur in the mouse transcriptome, and this is associated with resistance to injury-induced down-regulation, suggesting that the two motifs might function in a synergistic manner (Fig. S3C-D). To ensure that the observed associations are genuine, the same analysis was performed with the motifs on the 5’UTR or with random motifs. As expected, this control experiment does not show any significant association (Fig. S4). Taken together, the data suggest that CPE confers transcript stability against the global decrease induced by spinal cord injury by increasing RNA stability in conjunction with ARE motifs.

Although the 3’UTR-tested motifs are associated with higher resistance to injury-induced down-regulation, only CPE maintained this association in transcripts included in CNS development, axon development, and cell morphogenesis GO categories. In addition, CPE-containing genes include validated promoters of axonal regeneration such as Cebpβ and c-Jun (Fontana et al., 2012; Yan et al., 2009), which are among the CPE containing transcripts with the highest upregulation upon injury in the total RNA fraction (Table S1). Together with the fact that Cpeb1 overexpression in *Drosophila* promoted robust axonal outgrowth of developing sLNvs (Table 1), we chose Cpeb1 for further detailed investigation.

To elucidate whether CPE has a general functional role, GO enrichment studies were performed on the prevalence of CPE among all protein coding genes of the mouse and fly genomes. Many nervous system development categories were enriched among CPE containing genes in the mouse genome, including neuron projection morphogenesis, axonogenesis and axon guidance (Fig. 3C and Table S5). There were no categories found with an under-representation of CPE. Interestingly, almost all categories in the mouse genome enriched in CPE-containing genes are also enriched in *Drosophila*, suggesting a high level of conservation of CPE function between the two species.

### Cpeb1 promotes regeneration following neuronal injury

Our data suggest that CPE-enriched transcripts are temporarily protected from degradation after injury. Surprisingly, neither Cpeb1 mRNA nor protein levels were changed significantly upon SCI (although a trend to down-regulation could be observed (Fig. S5). Whether Cpeb1 is specifically down-regulated in neurons following SCI could not be assessed due to the lack of a working anti-Cpeb1 antibody for immunohistochemistry. To address the neuronal specific role of CPEB1 in axonal regeneration, and specifically whether the failure to upregulate Cpeb1 might in part explain the transient and abortive nature of the regenerative response to injury, we overexpressed the fly homologue of Cpeb1, Orb, exclusively in the sLNvs. Fly brains were dissected and kept in culture as described (Ayaz et al., 2008; Koch and Hassan, 2012) (see also accompanying manuscript). sLNvs were mechanically injured and the regenerative response was assessed after four days. The number of sprouts, regenerated length and distance reached from the lesion point were found to be significantly increased in neurons overexpressing Orb compared to control flies overexpressing LacZ in the same set of neurons (Fig. 4A-D).

**Figure 4:**
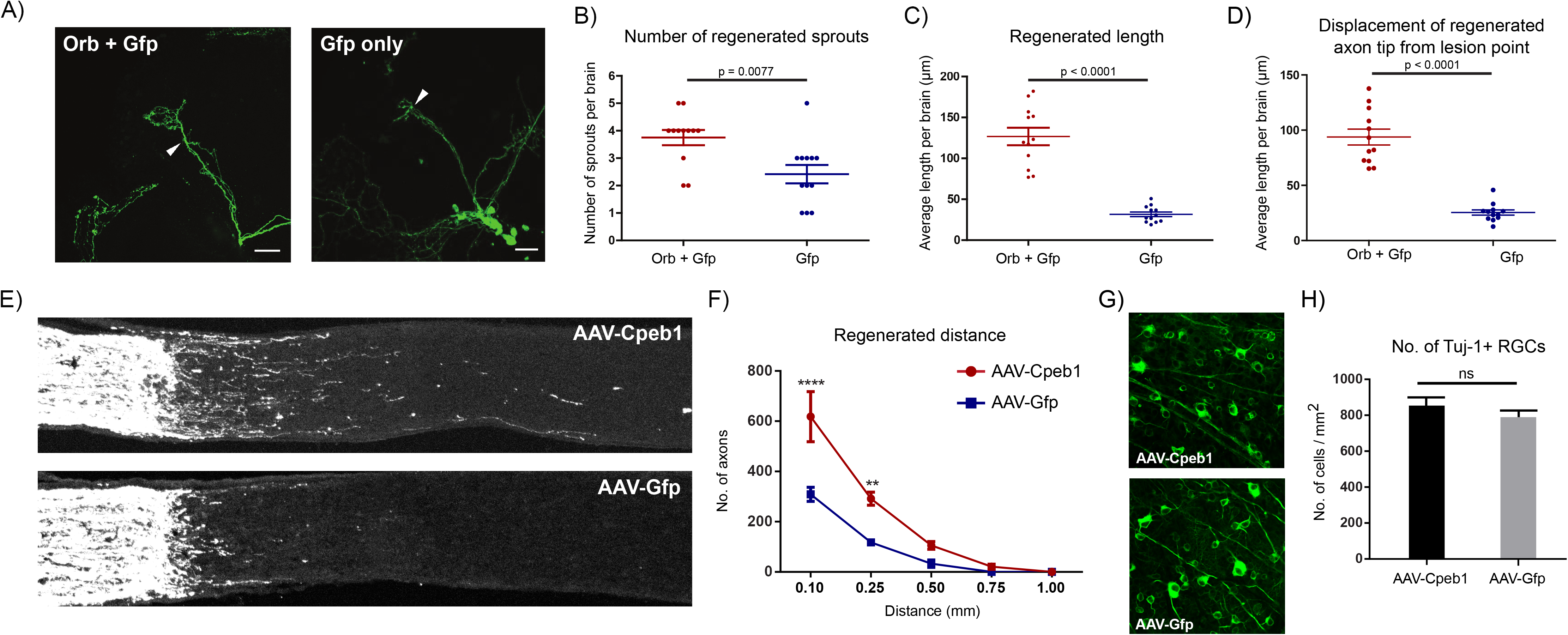
Cpeb1 overexpression promotes axonal regeneration in the adult mouse and Drosophila CNS. A-D) Over-expression of Orb (Cpeb1 homolog) enhances axonal regeneration in Drosophila sLNv neurons 4 days after axotomy. A) Representative images. Arrowheads indicate lesion points. Scale bars: 30μm. B-D) Quantification of number and length of regenerated axon sprouts. Each point represents one brain slice from one fly. n=13 (Orb+Gfp) or 12 (Gfp only) flies. Error bars: mean +/- S.E.M. E-H) Over-expression of Cpeb1 enhances mouse axonal regeneration 2 weeks after a optic nerve crush injury without affecting RGC survival. E-F) Representative images and quantification of traced axons following crush injury. G-H) Shows representative images and quantification of whole-mount staining of Tuj1+ cells in the retina.

To investigate whether this effect is conserved in mammals too, we turned to a mouse model of optic crush injury that allows overexpression of Cpeb1 in mouse retinal ganglion cells (RGCs) via infection with adeno-associated viral (AAV) vectors. Thereafter, regeneration of RGC-axons was assessed following a crush injury of the optic nerve. Importantly, as in *Drosophila*, over-expression of Cpeb1 enhanced axon regeneration in the mouse optic nerve, as both the number and length of regenerated axons were higher when measured 2 weeks after injury as compared to AAV-GFP infected control RGCs (Fig. 5E-F). The number of RGCs in the retina remained constant, indicating that the regenerative effect is not due to reduced cell death after injury (Fig. 5G-H). To test whether knockout of Cpeb1 produces an opposite effect, we knocked out Cpeb1 in primary cultures of mouse cortical neurons. Efficient knockout is triggered via AAV-Cre mediated deletion of exon 4 (Fig. S6A), which causes a frameshift that affects the activation and RNA recognition domains of Cpeb1. Neurons were cultured in a transwell chamber which specifically allows neurites to grow on the underside of the chamber. Scraping the lower side of the transwell mimicked a transection-injury. Thereafter, regenerating neurite on the underside could be examined. Notably, knockout of Cpeb1 was found to reduce the number and length of regenerated neurites 24 hours after injury (Fig. S6B-C). Together, these data support the notion that Cpeb1 is an enhancer of regeneration, and that this function is conserved between mice and *Drosophila*.

## DISCUSSION

We profiled the responses to spinal cord injury both at transcriptome and translatome level, and find them to be highly uncoupled. A screening of factors showing this uncoupled behaviour in *Drosophila* sLNvs revealed that 50% of the transcripts being prioritized for translation despite exhibiting reducing levels following spinal injury modulated axonal growth of developing neurons in the fly. Further, in silico analysis of these uncoupled-transcript led to identification of the CPE motif as highly conserved across species in functions including CNS-development. To influence expression of CPE-containing transcripts we overexpress Cpeb1 in injured Drosophila and mammalian neurons. CPEB1 emerged as a conserved necessary and sufficient activator of neuronal regeneration. The current approach also illustrates the feasibility of uncovering novel functions of RNA-regulatory proteins by combining studies in mammalian CNS with screenings in model organism, such as Drosophila.

The substantial uncoupling between transcription and translation in the injured spinal cord highlights the importance of post-transcriptional gene regulation in axon regeneration. The finding that the majority of genes are decreased in polysome-bound RNA agrees with previous observations that stress conditions trigger a general shut down of translation to maximize cell survival (Park et al., 2008; Yamasaki and Anderson, 2008). The use of the uncoupled response in RNA regulation as a selection criteria proved to be more efficient at discovering novel factors influencing axonal growth than selection based on prior knowledge on their role in neural and neurite development. In addition, the fact that Cpeb1 has a positive role in axonal regeneration and the similar enrichment of CPE in the genomes of both mouse and *Drosophila*, indicates that the role of Cpeb1 is conserved across species.

These findings agree well with the increasing number of studies that report the role of axonal translation of specific mRNAs for axonal regeneration (Hanz et al., 2003; Perlson et al., 2005; Perry et al., 2012; Yan et al., 2009; Yudin et al., 2008). In order for localized translation to occur in axons, both mRNA, as well as translation machinery, such as ribosomes and translation factors (e.g. elF4E), need to be present. In the case of mRNAs, it was found that injury triggers substantial changes in the axonal mRNA repertoire in cortical neuron cultures (Taylor et al., 2009). In an *in vivo* setting, growth cone-associated mRNAs Gap-43, Nrn1 and ActB are increased in crush-injured regenerating sciatic nerve axons compared to naïve ones (Kalinski et al., 2015). The same study also reported that the presence of ribosome components and activated translation factors in sciatic nerve axons is induced by injury. A preconditioning injury that activates regeneration of secondarily injured DRG axons increases the rate of incorporation of radioactively labeled amino acids, and selective local axonal application of the translation inhibitor cycloheximide severely reduces regenerative response (Verma et al., 2005). Together, this confirms that axonal translation occurs upon injury. Remarkably, the levels of ribosome components, translation factors and growth cone-associated mRNAs in regenerated transected spinal cord axons were comparable to those of crush-injured regenerating sciatic axons. This suggests that increasing axonal protein synthesis of specific mRNAs may just be key to overcome the lack of regeneration in the CNS (Kalinski et al., 2015).

While we identified Cpeb1 as an enhancer of axonal regeneration and CPE to be associated with expression changes upon spinal cord injury, it is surprising that this occurs in total RNA and not polysome-bound RNA, since Cpeb1 is best known for its role in cytoplasmic polyadenylation and translation control. However, Cpeb1 is also associated with a myriad of functions in post-transcriptional regulation, including alternative polyadenylation, RNA transport, storage and degradation. Cpeb1 has been shown to transport CPE-containing mRNAs into dendrites in rat hippocampal neurons, in particular Map2, as ribonucleoprotein (RNP) and in a microtubule-dependent manner (Huang et al., 2003). In addition, Cpeb1 is present in stress granules and dcp1 bodies, which are subcellular structures for mRNA storage and degradation (Anderson and Kedersha, 2006; Eulalio et al., 2007). Overexpression of Cpeb1 increases the assembly of these structures and is dependent on its RNA binding domain (Wilczynska et al., 2005). Interestingly, this is not dependent on the phosphorylation site for activation of Cpeb1 during cytoplasmic polyadenylation. Similarly, deletion of the phosphorylation site does not alter the distribution of Cpeb1-containing foci within the dendrites and synapses of Xenopus optic tectal neurons, suggesting that the function of Cpeb1 to transport and target mRNAs to RNP complexes is independent of its role in cytoplasmic polyadenylation (Bestman and Cline, 2009). Foci containing inactive Cpeb1 mutants located near synapses do, however, show higher intensities than those containing wild-type Cpeb1, suggesting that inability to activate translation may trap Cpeb1 and its target mRNA within the RNP complex (Bestman and Cline, 2009). A possible scenario linking these observations is that inactive Cpeb1 forms RNP complexes together with the bound mRNA, leading to simultaneous repression of mRNA translation and protection from degradation while guiding them to the final location. Another scenario involves regulation of alternative polyadenylation by Cpeb1, thereby recruiting splicing factors to different polyadenylation sites and generating transcripts of varying 3’UTR lengths (Bava et al., 2013). This process affects the cis-elements present in the 3’UTR of the transcript (e.g. other RNA binding motifs or miRNA binding sites), which in turn affects localization, stability and translation efficiency of the transcript (Lianoglou et al., 2013; Mayr and Bartel, 2009).

In summary, starting with a global approach our study revealed the role of wide-spread post-transcriptional regulation in the early injury response in the spinal cord. While translation of mRNAs related to CNS development appears to be prioritized in the acute phase after injury, limitation in their transcript availability likely leads to the eventual failure in regeneration. By focusing on genes that exhibit uncoupling behavior between transcript availability and translation, we identified a number of genes that modulate outgrowth of axons during development, demonstrating that this as a viable method to identify neuronal intrinsic regulators of regeneration. We have also found the association of 3’UTR motifs with CNS injury response, and identified Cpeb1 as a modulator of axon regrowth, possibly by increasing the availability of development-related transcripts.

## METHODS

### Mouse experiments

#### Spinal cord injury

Female C57BL/N mice of 10 weeks of age were used for the spinal cord injury. Animals were bred in house at the DKFZ Center for Preclinical Research Facility and housed under standard conditions. Animals were subjected to laminectomy and then an 80% transectional spinal cord injury by cutting the spinal cord with irridectomy scissors. Naïve mice were subjected only to laminectomy. Nine hours after the operation, 2.5 cm of spinal cord tissue centering on the lesion site was extracted. Procedures were conducted in accordance with the DKFZ guidelines and approved by the Regierungpräsidium Karlsruhe.

#### Translation state array analysis (TSAA)

RNA was isolated from the chosen tissue and segregated into a total RNA fraction and a polysome-bound RNA fraction. Polysome-bound RNA was isolated via fractionation in sucrose gradient as previously described (Lou et al., 2014), with fractions containing RNA bound to two or more ribosomes collected.

The samples were profiled with Affymetrix arrays (model Mouse430_a2) with RNA input amounts of 5 µg and 3µg for total and polysome-bound RNA respectively. <u>Array data is accessible from GEO (GSE92657).</u> Array data corresponding to each fraction were normalized separately, as polysome-bound RNA is a subset of total RNA and standard microarray normalization methods that assume equality of distributions of total intensity between arrays could not be used.

Normalization was performed using the vsn method as implemented in the vsnrma function from the R/bioconductor package vsn (Huber et al., 2002). lts.quantile=0.5 was used to allow for robust normalization when many genes are differentially expressed. Differential expression was calculated using limma (Smyth, 2004) from R/bioconductor at the level of probesets, the parameter “trend” was set to TRUE for the empirical moderation of standard errors. Probesets with Benjamini-Hochberg false-discovery rates (FDR) <0.05 were considered as differentially expressed. Probesets were then translated to Ensembl gene IDs (Ensembl v72; www.ensembl.org), and those mapping to multiple IDs were excluded from subsequent analysis. For cell marker analysis normalised microarray intensities were averaged over replicates. The expression values for 103 measured marker genes were used to show the composition of samples.

#### Gene Ontology enrichment

Enrichment analysis of GO biological process categories between up- and down-regulated genes were performed separately for each of the RNA fractions. To this end, up-regulated genes were compared against a background of all differentially expressed genes with the hypergeometric distribution, using GOstats (Falcon and Gentleman, 2007) and annotation from the org.Mm.eg.db v2.14.0 package, both from R/bioconductor. Under-representation of up-regulated genes in a category is equivalent to having an enrichment of down-regulated genes, and is displayed as such.

For the enrichment study of CPE in the mouse and fly genomes, GO annotations from annotation packages org.Mm.eg.db v2.14.0 and org.Dm.eg.db v2.14.0 with experimental evidence code (i.e. EXP, IDA, IPI, IMP, IGI, IEP) were used. The selectiveness is because GO annotations are based on different kinds of evidence including homology, and circularity would occur when comparing results from two different genomes with homology included. To prevent artificially inflating the number of motifs when genes have more than one transcript with common 3'UTR, we performed the enrichment analysis at the level of genes. Transcripts with annotation of transcript biotype as protein coding were translated to Ensembl gene ID and then to Entrez. Genes with only CPE-containing transcripts were compared against all genes (excluding those with both CPE-containing and CPE-free transcripts) with the hypergeometric distribution using GOstats.

Results from the enrichment analyses were visualized using Cytoscape v3.0.2 (Shannon et al., 2003). Results were represented on the GO subnetworks comprising of categories significant in at least one of the comparisons (Benjamini-Hochberg FDR <1e-4 for Fig. 2 and <1e-5 for Fig. 3C), with each node referring to one GO category. Color represents the direction of enrichment, and the intensity of color and size of the node represent significance (Benjamini-Hochberg BH FDR >0.05 is depicted as white).

#### UTR motif analysis

The UTR sequences from transcripts belonging to genes with Ensembl annotation as known and protein coding were obtained from Ensembl v72 and searched for regular expressions representing motifs. Sequences used for each motif (Piqué et al., 2008; Spasic et al., 2012; Tian et al., 2005) are listed in Table S4.

Probesets were translated to Ensembl transcript ID (Ensembl v72), and only probesets mapping a unique transcript were used. Distributions of log2 (fold change) were compared using the Kolmogorov-Smirnov test in R (http://cran.rproject.org/). To represent the expression change profiles for transcripts from specific GO categories, we used GO annotation at the level of transcript downloaded from Ensembl v72 with the R/Bioconductor package biomaRt.

#### Optic nerve crush injury

Experimental procedures were performed in compliance with animal protocols approved by the Animal and Plant Care Facility at the Hong Kong University of Science and Technology. C57BL/6 mice of 5-6 weeks of age were anesthetized with ketamine (80 mg/kg) and xylazine (10 mg/kg) and received Meloxicam (1 mg/kg) as analgesia after the surgery. AAV2 vectors expressing either Cpeb1 or Gfp under the neuron-specific human synapsin promoter were injected into the vitreous bodies with a Hamilton microsyringe. Five weeks after vector injection, the optic nerve was gently exposed intraorbitally and crushed with jeweler’s forceps (Dumont #5; Fine Science Tools) around 1 mm behind the optic disk. Mice were kept for 2 weeks after injury before tracing. To visualize RGC axons in the optic nerve, 1.5 μL cholera toxin β subunit conjugated with Alexa555 (CTB555, 2 μg/μl, Invitrogen) were injected into the vitreous bodies. Two days after the CTB injection, animals were sacrificed by transcardial perfusion for histology examination. In each mouse, the completeness of optic nerve crush was verified by showing that anterograde tracing did not reach the superior colliculi.

#### Immunofluorescence staining of retina and optic nerve and quantifications

Eyes and optic nerves were cryosectioned and examined under an epifluorescence (Nikon, TE2000) or confocal microscope (Zeiss, LSM Meta710). Total numbers of RGCs were determined by whole-mount retina staining. A mouse anti-Tuj1 antibody (Covance) was used to detect RGCs. Twelve images (3 for each quarter, covering peripheral to central regions of the retina) from each retina were captured under a confocal microscope (400X) and Tuj1-positive cells were counted in a blind fashon. To detect traced axons after optic nerve crush, longitudinal sections of optic nerves were serially collected. To analyze the extent of axonal regeneration in the optic nerve, the number of axons that passed through distance d from the lesion site was estimated using the following formula: ∑a_d_ = πr^2^ × [average axon numbers per mm/t], where r is equal to half the width of the nerve at the counting site, the average number of axons per mm is equal to the average of (axon number)/(nerve width) in 4 sections per animal, and t is equal to the section thickness (8 μm). Axons were manually counted in a blind fashion. Two-tailed Student’s t-test was used for the single comparison between two groups. The rest of the data were analyzed using ANOVA. *Post hoc* comparisons were carried out when a main effect showed statistical significance.

#### Generation and primary culture of Cpeb1 knockout neurons

Primary cultures of cortical neurons were prepared from embryos of Cpeb1^flox/flox^ mice. Cre-mediated excision will remove exon 4 of Cpeb1 and cause a frameshift that affects the phosphorylation site for activation of Cpeb1 as well as its RNA recognition motifs. Cultures were prepared from cortices of E16.5 embryos. Briefly, dissected cortices were digested with 0.05% trypsin for 15 minutes, triturated with a fire-polished glass pipette and plated on poly-L-lysine coated surfaces. Neurons were cultured in HS-MEM (1x MEM (Thermo Fisher), 10% horse serum, 1.2% glucose, 4mM L-glutamine, 1mM sodium pyruvate, 0.22% NaHCO_3_, 100 U/ml penicillin-streptomycin) and infected with serotype 2 AAV encoding CAG-Cre at an MOI of 1×10^5^. 24 hours later medium was replaced with N2B27-MEM (1× MEM (Thermo Fisher), 1× N2 supplement, 1× B27 supplement, 0.1% ovalbumin, 0.6% glucose, 2mM L-glutamine, 1mM sodium pyruvate, 0.22% NaHCO_3_, 100 U/ml penicillin-streptomycin).

#### Western blotting

Western blotting against Cpeb1 in spinal cord tissues was performed with standard procedures using an antibody raised in-house by the group of H. Zentgraf. Cell lysates of murine hippoccampal HT22 cells transiently over-expressing Cpeb1 were used as positive control.

#### *In vitro* regrowth assay

Neurons were cultured on Fluoroblok transwell chambers with PET membranes with 3µm pores (Millipore) that allow extension of neuronal processes to the underside of the membrane while keeping cell somas on the top. After 7 days in culture, the underside was scraped with sterile cotton swabs to cut the processes. 24 hours after injury, processes were labeled with addition of 1µM of Calcein AM (BD Biosciences) to the culture medium and imaged with a Zeiss Cell Observer epifluorescence microscope. Images were analyzed and traced with a custom wrote macro in ImageJ in a blind manner. A mixed effects model was used for statistical comparison to account for variability in various levels of the experimental setup.

### Drosophila experiments

#### Stocks and genetics

*Drosophila* melanogaster stocks were kept on standard cornmeal media. For tissue specific overexpression of the transgenes, we used the GAL4/UAS system (Brand and Perrimon, 1993). For the genetic screen in development and after injury, pdf-gal4, uasgfp; pdf-gal4, uas2x egfp/cyo flies were kept as a stock and used to drive expression of the various candidate genes, or crossed to wild-type Canton S (CS), in the case of the outgrowth experiments, or to UAS-lacZ, in the case of the injury experiments. Overexpression stocks were obtained from the Bloomington Stock Centre and the PDF-Gal4 line was obtained from P. Taghert. All flies were dissected 2-10 days after eclosion.

#### Drosophila outgrowth and injury assays

To measure axonal outgrowth during development, flies were reared at 25°C and were dissected in fresh PBS. A minimum of 5 fly brains (10 sLNv projections) per genotype were fixed in 4% formaldehyde and stained with an anti-GFP antibody (Molecular Probes; 1:500), according to standard methods (Ayaz et al., 2008). For the axonal regrowth analysis after injury, flies were reared at 18°C (to minimize developmental effects), and shifted to 25°C one day prior to injury. Whole brain explants on culture plate inserts were prepared as described (Ayaz et al., 2008; Koch and Hassan, 2012). In brief, culture plate inserts (Milipore) were coated with laminin and polylysine (BD Biosciences). Fly brains were carefully dissected out in a sterile Petri dish containing ice cold Schneider’s *Drosophila* Medium (GIBCO), and transferred to one culture plate insert containing culture medium (10 000 U/ml penicillin, 10 mg/ml streptomycin, 10% Fetal Bovine Serum and 10 µg/ml insulin in Schneider’s *Drosophila* Medium (GIBCO). sLNv axonal injury was performed using an ultrasonic microchisel controlled by a powered device (Eppendorf) as described (Ayaz et al., 2008; Koch and Hassan, 2012)and dishes were kept in a humidified incubator at 25°C. Four days later, cultured brains were fixed and immunohistochemical staining was performed as for freshly dissected samples.

#### Imaging, morphometric measurements and statistics

For the outgrowth experiments, brains were visualized under a fluorescent microscope equipped with a GFP filter and classified as having “increased outgrowth”, “reduced outgrowth” or “no observable effect” according to comparison of sLNv length with that of controls.

For the injury experiments, de novo growth was assessed four days after injury by measuring injured sLNv projection that has formed at least one new axonal sprout of a minimum length of 12µm. The exact injury location was accessed by comparison with axonal projection length at 5 hours (where no de novo growth has occurred). Imaging was performed on a Zeiss 500 or 700 confocal microscope and analyzed with Image J. Regenerated length was defined as the de novo axon lengths using the manual tracing tool. Projected distance was defined as the displacement of the axon sprouts from the lesion site, measured in a straight line. All images were analyzed in a blind manner. Statistical comparisons were performed with two-tailed student’s t-test.

## ACKNOWLEDGMENTS

We thank Thomas Hielscher, Axel Benner, Tim Holland-Letz and Simone Braun from the DKFZ Biostatistics Division for statistical advice; Ha Nati from the University of Heidelberg for help with Cytoscape; and Damir Krunic from the DKFZ Light Microscope Core Facility for providing the macro for neuronal process tracing. This study was supported by the German Ministry of Education and Research (BMBF), the German Cancer Research Center (DKFZ), the Cluster of Excellence CellNetworks (EXC81), the German Research Foundation (DFG), the Helmholtz Initiative for Synthetic Biology, and the Hong Kong Research Grants Council 689913, 16101414.

## AUTHOR CONTRIBUTIONS

WPL performed validation experiments and wrote the manuscript. AM performed bioinformatics analysis and wrote the manuscript. MK and MZ performed the Drosophila experiments. MG performed bioinformatics analysis. SL and SK performed experiments. ES and DG generated the AAV vectors. RM and CM provided the transgenic mouse line, antibodies and critical discussion of the data. CY and NL performed the optic nerve crush injury experiments supervised by KL who also wrote the manuscript. BH supervised the Drosophila part of the study and wrote the manuscript. AMV designed and supervised the study and wrote the manuscript.

---

**Figure S1:**
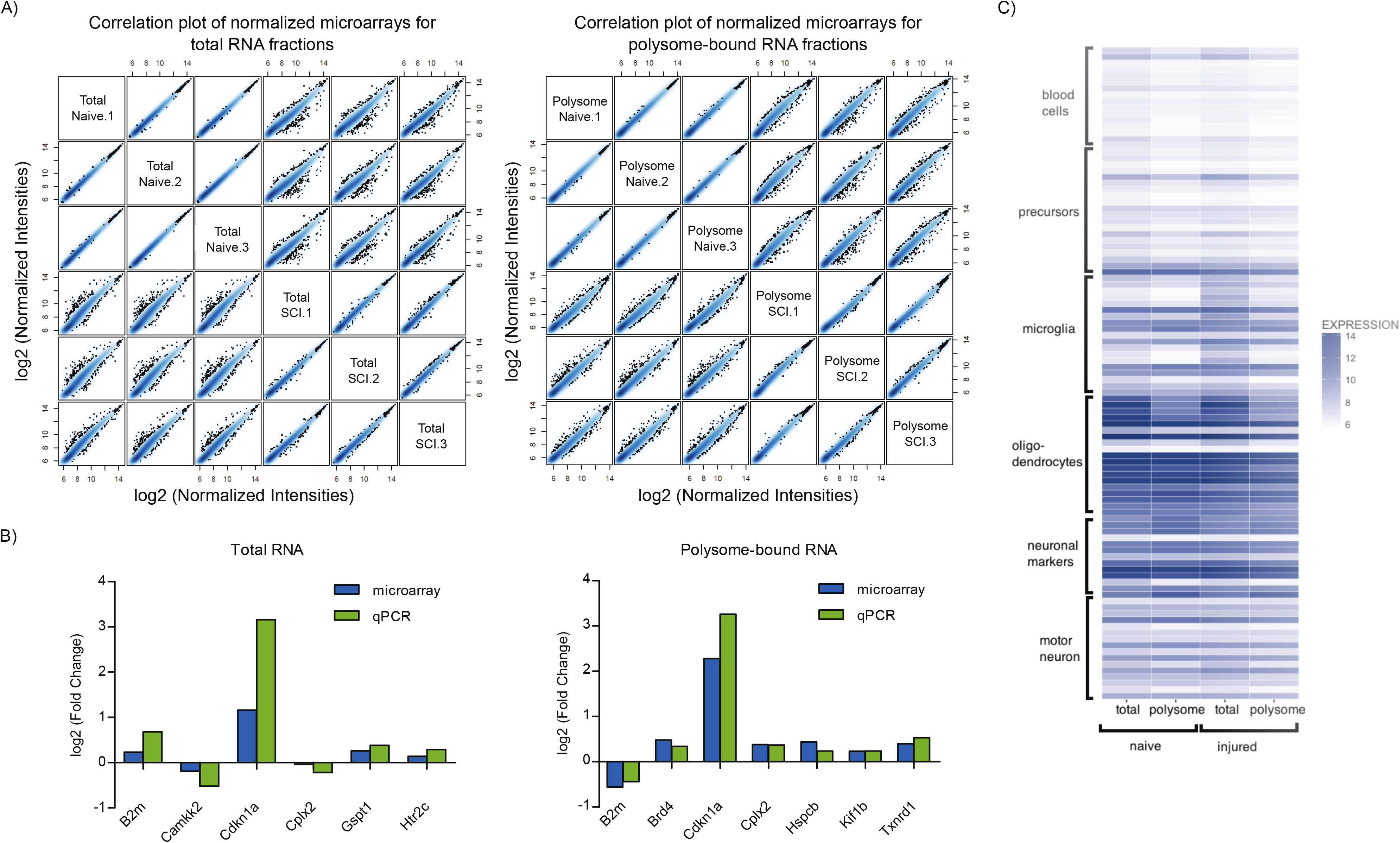
Correlation and validation of microarray. A) Correlation plot of normalised arrays (log2[normalised intensities]) for total and polysome-bound RNA fractions. B) Comparison of expression changes of selected genes derived from microarray or qPCR. C) Expression patterns of cell-type-specific genes are similar in both, the condition ‘naive’ and ‘injured’. Colours represent normalised intensities of microarray probes mapped to marker genes for motor neurones, other neurones, oligodendrocytes, microglia, precursors, and blood cells. The expression patterns between the conditions ‘naive’ and ‘injured’ of these markers show a Pearsons’ correlation of 0.97 for total RNA and 0.99 for RNA bound to polysomes.

**Figure S2:**
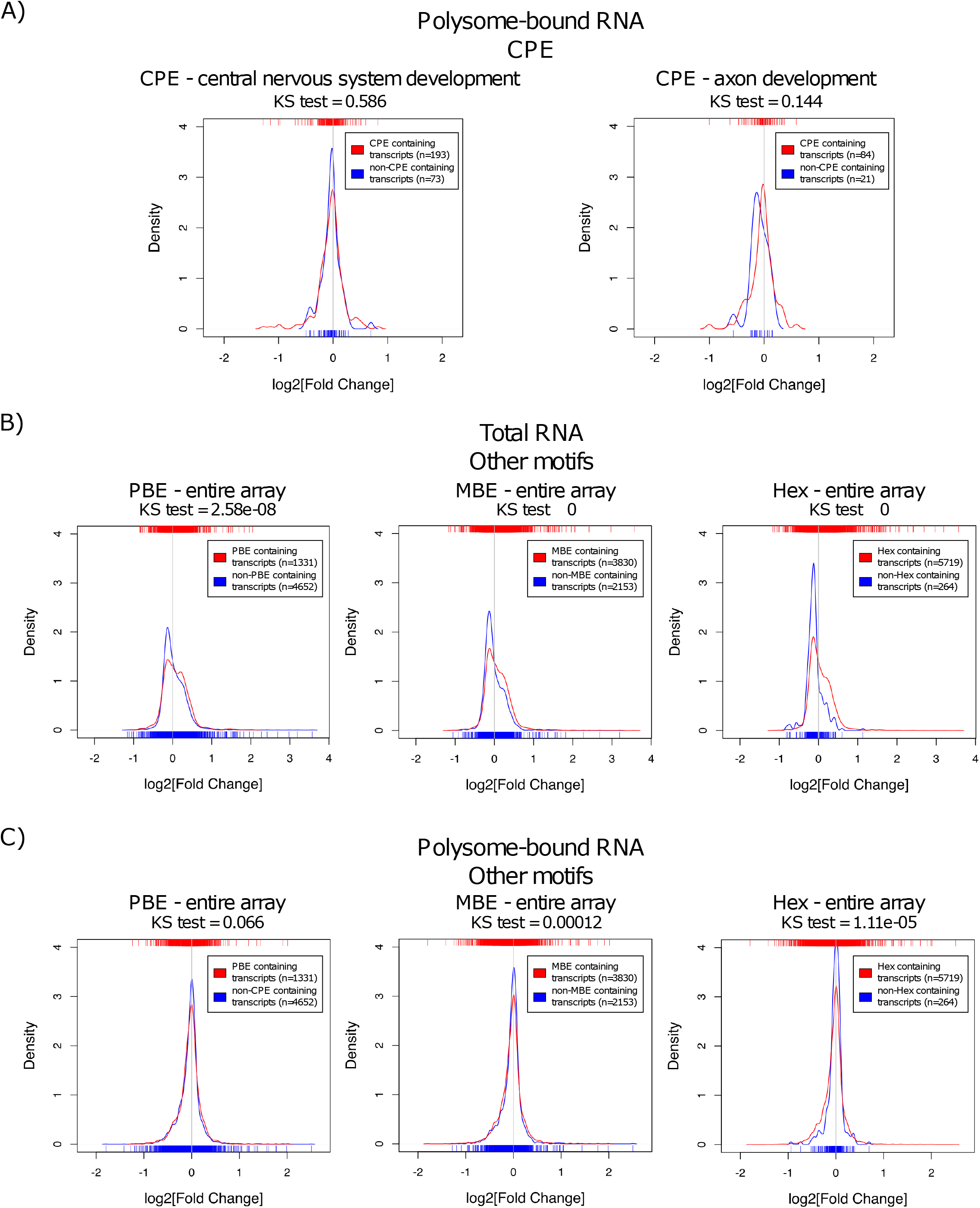
Association of other RNA motifs with expression changes upon SCI. A) Density curves showing distribution of expression changes in polysome-bound RNA upon SCI of transcripts containing CPE in the 3’ UTR. B-C) Density curves of expression changes in total and polysome-bound RNA upon SCI of transcripts separated by the presence of PBE, MBE and Hex in the 3’ UTR. Ticks on top and below the plots represent values of log2 (fold change) of individual transcripts. Distributions were compared with Kolmogorov-Smirnov test.

**Figure S3:**
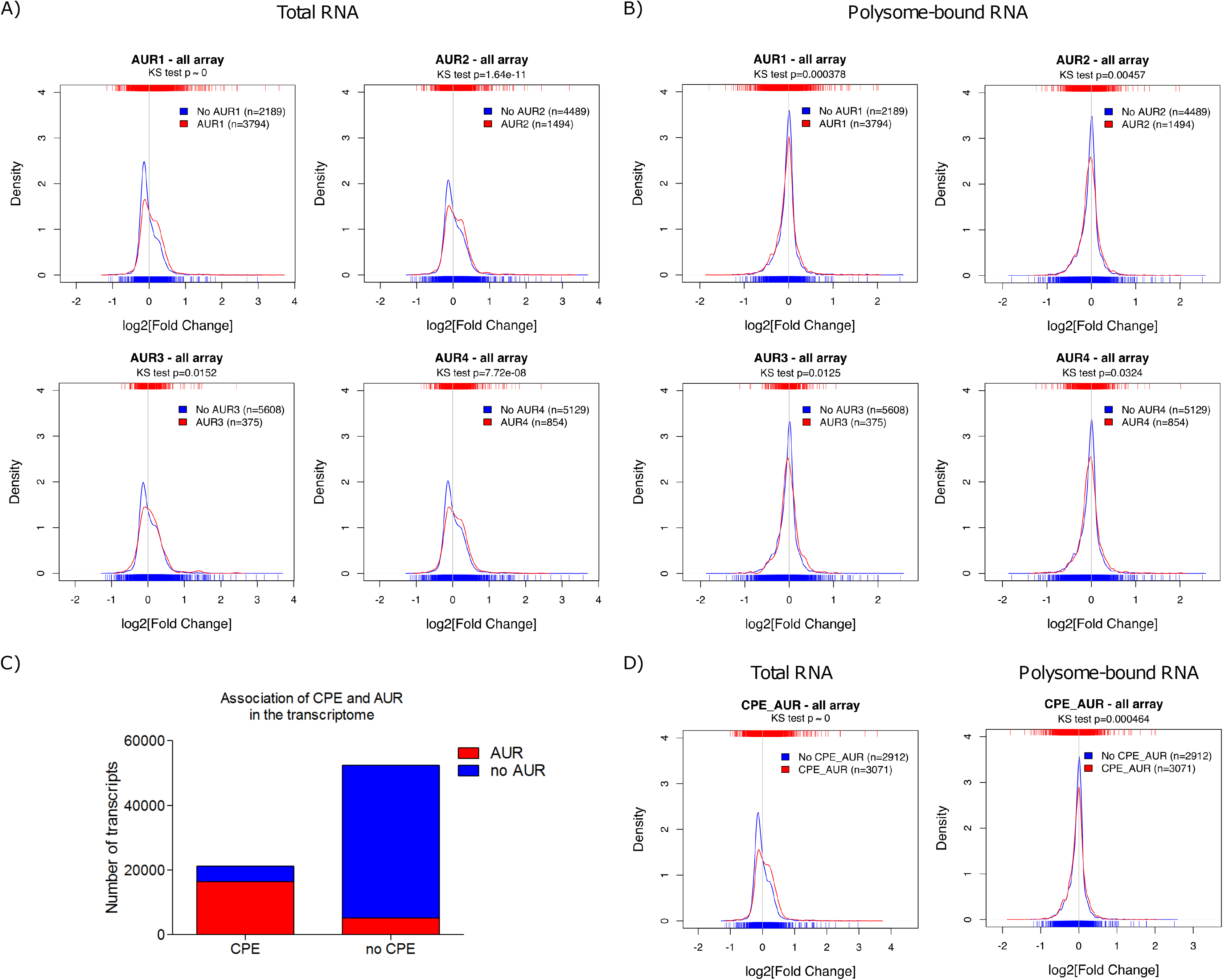
Association of AREs with higher expression changes upon SCI and with CPE. A-B) Density curves showing distribution of expression changes in total and polysome-bound RNA upon SCI of transcripts containing AREs in the 3’ UTR. C) Number of transcripts harbouring CPE and AUR in the whole transcriptome. D) Density curves showing the distribution of expression changes of transcripts containing both AREs and CPE in the 3'UTR in total and polysome-bound RNA upon SCI. Ticks on top and below the plots represent values of log2 (fold change) of individual transcripts. Distributions were compared with Kolmogorov-Smirnov test.

**Figure S4:**
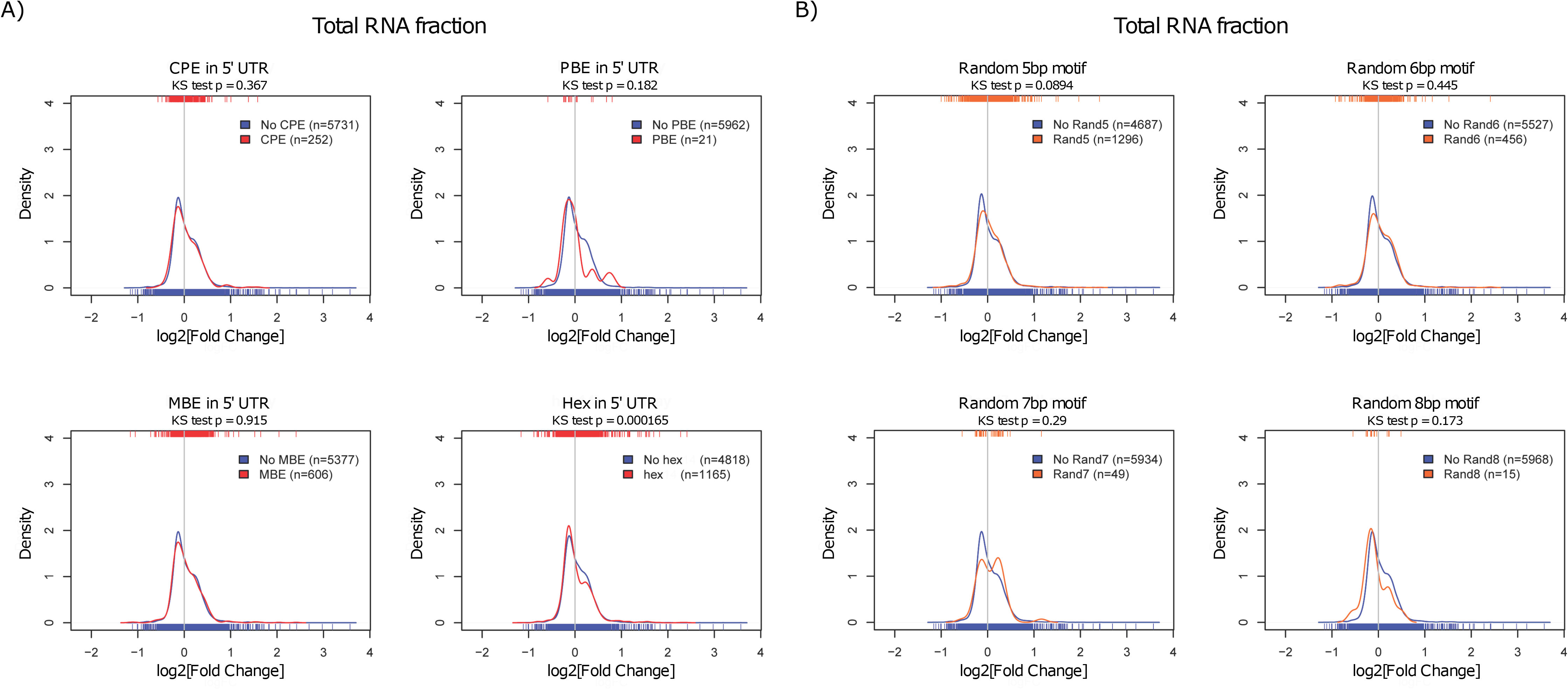
Control analysis for motif analysis. Substituting the motif analysis with A) the same motifs but in the 5’ UTR and B) random motifs shows no association with expression changes.

**Figure S5:**
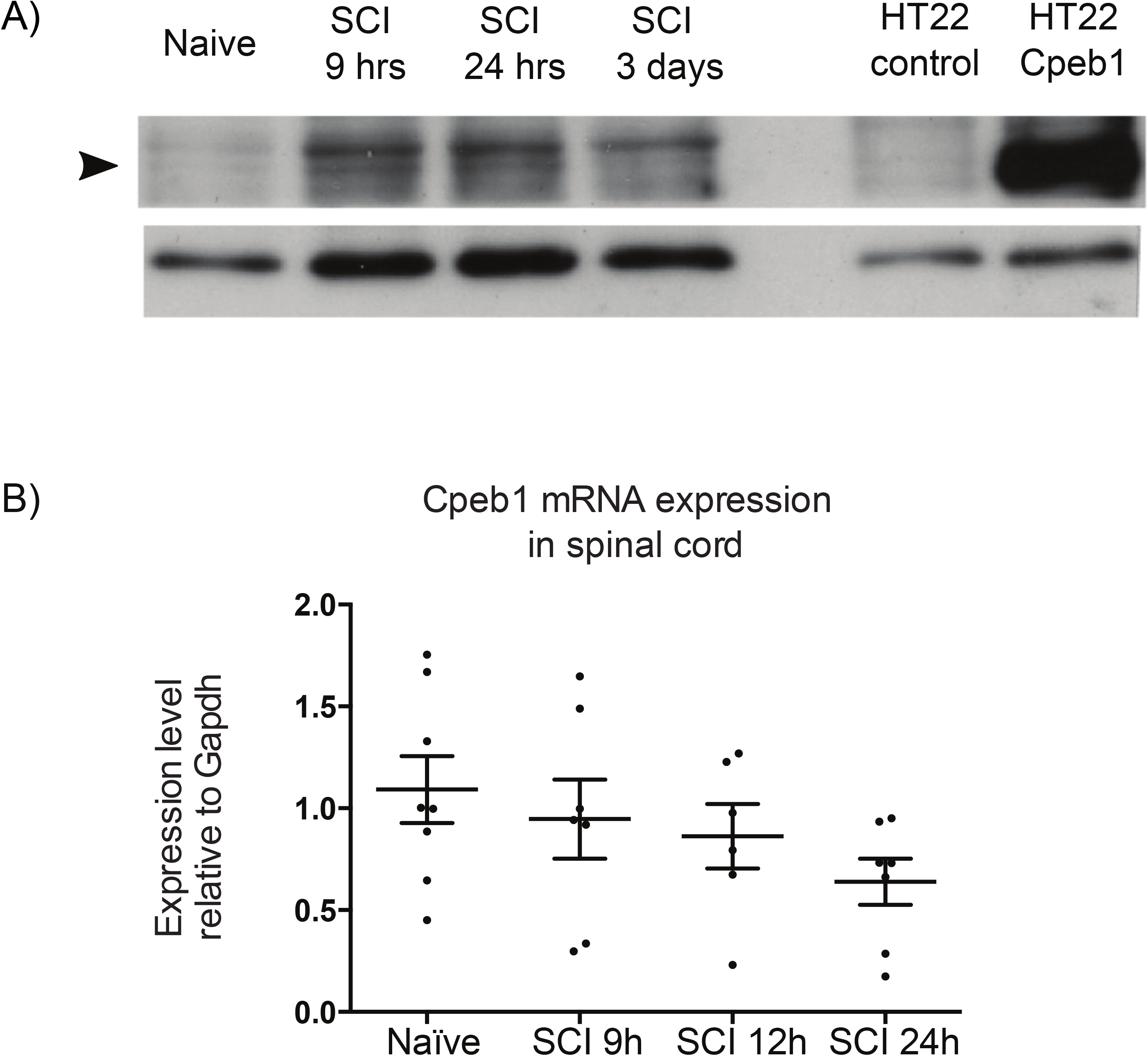
Expression of Cpeb1 detected in injured spinal cords. A) Western blotting of Cpeb1 of naïve and injured spinal cords. Cell lysates of HT22 cells transiently over-expressing Cpeb1 were used as positive control. B) qPCR of naïve and injured spinal cords for Cpeb1. (1-way ANOVA p=0.9622)

**Figure S6:**
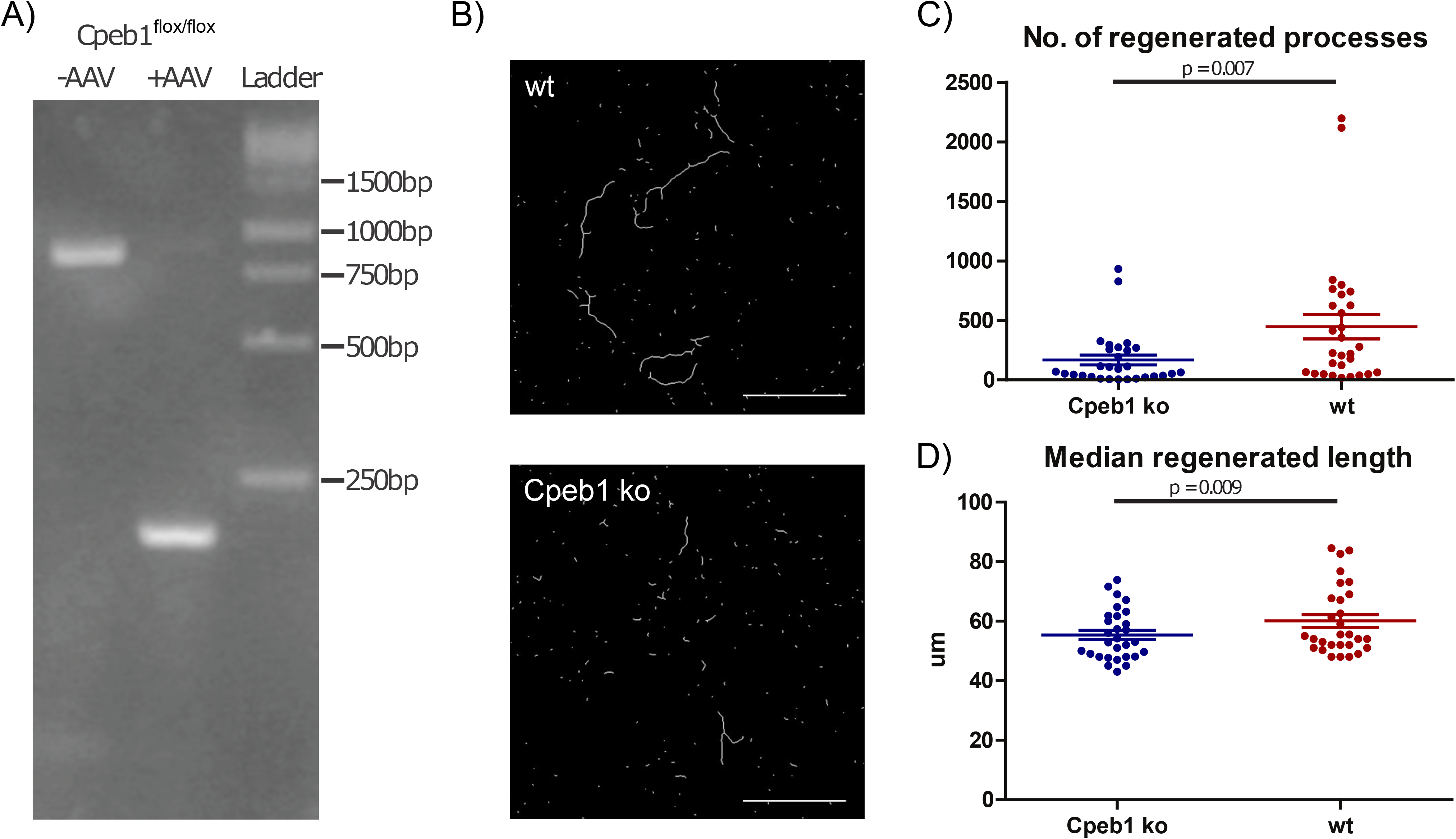
Deletion of Cpeb1 reduces neurite regeneration in vitro. A) Efficient deletion of Cpeb1 by AAV-delivered Cre confirmed by PCR. Expected sizes: 884bp (without deletion), 186bp (with deletion). B) Cortical neurons infected with AAV-Cre were seeded on transwell chambers, which exclusively allow neurite growth on the lower side of the membrane. Representative images of neurites skeletonized from image processing. C-D) Quantification of regenerating neurites 24 hours after injury. Each data point represents one culture chamber. A total of 29 culture chambers prepared from 4 mice were used per group. Cpeb1 ko: Cpeb1flox/flox + AAV; wt: wild-type + AAV. Scale bars: 100μm. Error bars: mean +/- S.E.M.

